# Structural insights into ATP-sensitive potassium channel mechanics: a role of intrinsically disordered regions

**DOI:** 10.1101/2022.08.03.502592

**Authors:** Katarzyna Walczewska-Szewc, Wiesław Nowak

**Affiliations:** Institute of Physics, Faculty of Physics, Astronomy and Informatics, Nicolaus Copernicus University in Toruń, ul. Grudziądzka 5, 87-100 Toruń, Poland

## Abstract

Commonly used techniques, such as CryoEM or Xray, are not able to capture the structural reorganizations of disordered regions of proteins (IDR), therefore it is difficult to assess their functions in proteins based exclusively on experiments. To fill this gap, we used computational molecular dynamics simulations methods to capture IDR dynamics and trace biological function-related interactions in the Kir6.2/SUR1 potassium channel. This ATP-sensitive octameric complex, one of the critical elements in the insulin secretion process in human pancreatic β-cells, has four to five large, disordered fragments. Using unique MD simulations of the full Kir6.2/SUR1 channel complex, we present an in-depth analysis of the dynamics of the disordered regions and discuss the possible functions they could have in this system. Our MD results confirmed the crucial role of the N-terminus of the Kir6.2 fragment and the L0-loop of the SUR1 protein in the transfer of mechanical signals between domains that trigger insulin release. Moreover, we show that the presence of IDRs affects natural ligands binding. Our research takes us one step further towards understanding the action of this vital complex.

## 1. Introduction

Over decades, it has been known that the paradigm linking a protein’s biological function to its well-defined three-dimensional structure is not always fulfilled. Most of well-known proteins still fold quite precisely, adopting their function-related structures. However, a significant proportion of proteins, or protein fragments, escape this rule and perform important tasks in the organism without exhibiting a defined secondary structure^1, 2^

Intrinsically disordered proteins (IDPs) or intrinsically disordered protein regions (IDRs) can adopt a whole ensemble of conformations and act while remaining unstructured. Analyses show that up to 67% of eukaryotic proteins contain at least one disordered region^1^. Those include the ATP-sensitive potassium (KATP) channels discussed in our paper.

Human KATPs are octameric transmembrane complexes commonly present in smooth muscles, cardiac myocytes, brain and pancreatic beta-cells^3^. Pancreatic KATP channels, main objects studied here, open or close in response to physiological changes in the ratio of ATP to ADP present in the cytosol. Their particular conformational states affect the concentrations of potassium ions on both sides of the membrane and through the resulting change in the membrane potential Kir6.2/Sur1 channels mediate insulin secretion from pancreatic beta-cells. Dysfunctions of the channel caused by mutations may lead to neonatal diabetes^4^ or congenital hyperinsulinism^5^.

IDRs are commonly found in transmembrane proteins, regulating their permeation and recruitment of downstream interaction partners^6, 7^. Recent CryoEM studies indicate that pancreatic human KATPs have at least five regions which may be classified as IDRs. One may expect those regions to play a role in the KATP function. However, they were not analyzed so far. Particularly, this is the case for 32 amino acids long, unstructured, the N-terminus of the Kir6.2 protein (N-ter), whose relevance in the channel function was indicated by many experimental studies^8, 9^. Besides a glutamate-rich linker of SUR was suggested recently by Sung et al^10^ as a component of a conformational pathway toward vascular KATP channel activation. In multi-domain systems, IDRs often mediate the propagation of a perturbation arising at a specific location in one domain to a distal location^2^. Therefore, given that the cytosolic part’s interface between KATP domains is built mainly from IDRs (Figure 1b), this issue is worth investigating.

**Figure 1.**
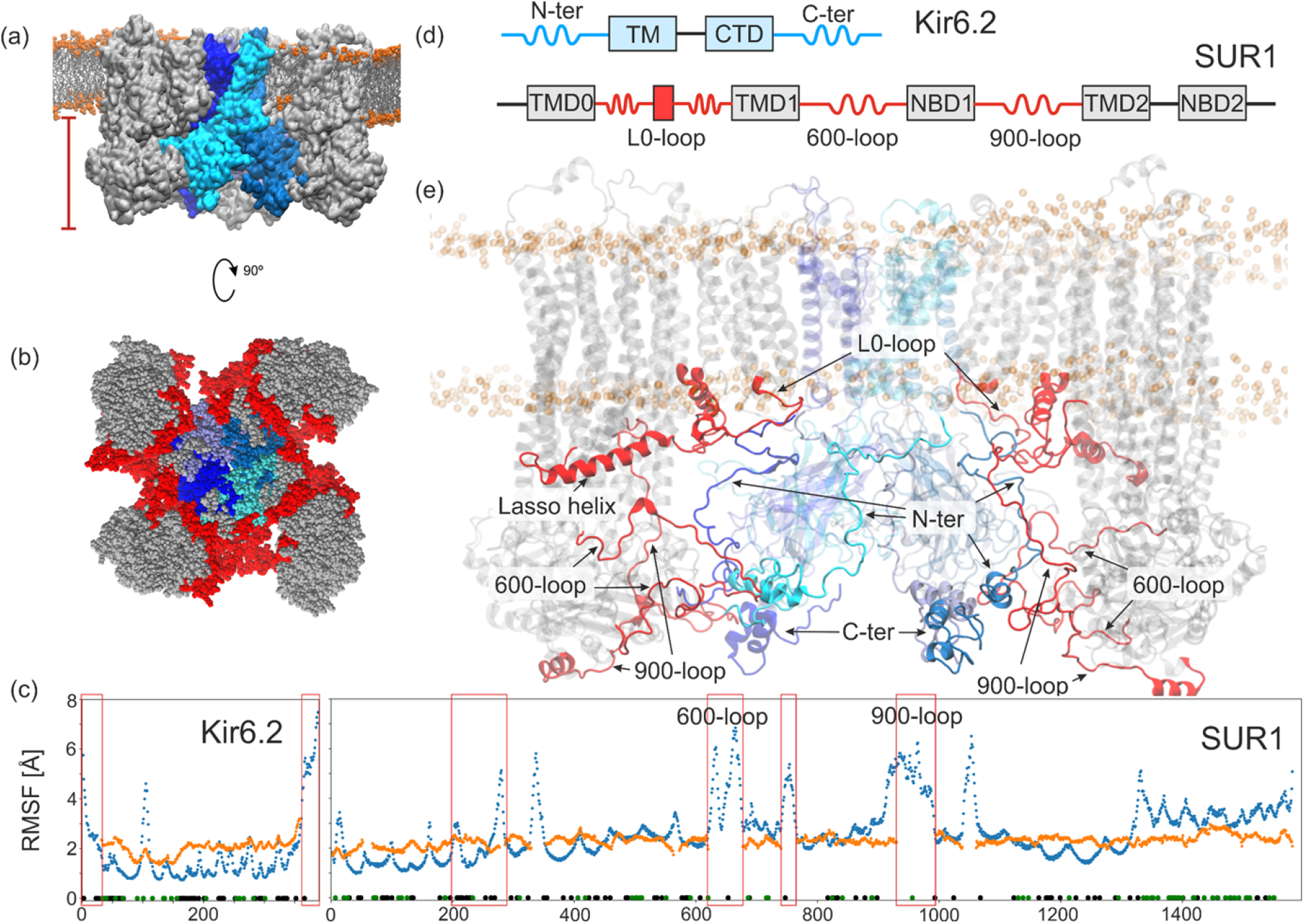
Simulated system of KATP octamer (a). SUR1 are in grey, Kir6.2 subunits are in shades of blue, lipid heads phosphorus atoms are orange. The cytoplasmic part of the KATP complex with the inter-domain interface formed purely with IDRs (red) (b). RMSF of Kir6.2 and SUR1 structures during MD simulation (blue) compared to an experimental temperature factor (orange)^12^ (c). The domain structure (d) and the cross-section of the channel (e). Intrinsically disordered regions (N-ter, C-ter, 600-loop, and 900-loop) and partially disordered region (L0-loop) are indicated in red and blue for SUR1 and Kir6.2, respectively.

In this paper, using novel, based on CryoEM data, models of the human KATP channel and the first all-atom molecular dynamics simulations of the complete KATP system on microsecond scales, we explore conformational space of IDRs in KATP. We monitor and show how IDRs structures evolve over time. We check whether they adopt any transient, well defined, conformations or yet they remain completely disordered on the present simulations timescale. We also discuss possible roles of IDR as mediators of interactions between KATP subunits. <u>We investigate how IDRs may affect the dynamics of the protein in the form we know so far</u>. Such knowledge will help us to understand better how KATP works.

## 2. Results and discussion

### 2.1. Where IDRs in KATP are located? Overview of major IDRs in KATP

The KATP complex consists of eight units (see Figure 1 a, b). Four of them, named Kir6.2, form the inner part lining the inward-rectifier potassium channel’s pore. Outside this inner core are located four sulfonylurea receptors - SUR1 units. They have an important regulatory function. SUR1 mediates the activation effect of Mg-ADP and affects the pharmacological profile of the channel^11^. The structure of the main channel components is well known^12–14^.

Both Kir6.2 and SUR1 subunits consist of transmembrane and cytosolic parts. In the case of Kir6.2 subunits, there are two transmembrane helices (TM) and one cytoplasmic domain (CTD). SUR1 has the structure consistent with other ABC transporters architecture - it is composed of two membrane-spanning domains (TMD1 and TMD2), two cytosolic nucleotide-binding domains (NBD1 and NBD2) and an additional transmembrane domain (TMD0) functioning as an interface between Kir6.2 and SUR1. In the whole KATP complex, 15% of the structure is not defined in the available CryoEM data^12^. These include 17% of Kir6.2 and 14% of SUR1. Most of such regions are intrinsically disordered (IDR-intrinsically disordered regions, see Figure 1c-e). It is assumed that the dynamical nature of IDRs makes them “invisible” for structural characterization methods like X-ray crystallography or Cryo-EM^15, 16^.

In Kir6.2, the major IDRs are N-terminus (**N-ter**) and C-terminus (**C-ter**) which partially form the cytoplasmic domain. In SUR1, three loops: the **600-loop** connecting TMD1 to NBD1, the **900-loop** connecting NBD1 to TMD2, and the **700-loop** belonging to NBD1 are fully disordered in known CryoEM structures. These fragments meet the accepted definition of an IDR, i.e. a fragment of 25 or more residues without a noticeable secondary structure^1^.

Moreover, polar (S, T, Q, D, E, R, K, H) and structure-breaking (G,P) residues are more common in such regions and hydrophobic, aromatic, and cysteine and asparagine residues are rare. Here we have included fragments located on the inner side of the cell membrane

We also decided to analyze the L0-loop with Lasso helix motif connecting TMD0 to TMD1 which is partially disordered (although nearly fully covered in CryoEM data), because of the potentially important role this fragment plays in the KATP function.

Below we provide a short description of each selected, disordered fragment.

The N-terminal end of Kir6.2 (**N-ter**) is recently a subject of vigorous research, as it was postulated that this flexible polypeptide chain has a major factor in controlling KATP opening/closing processes^8, 9^. Our previous studies have confirmed that N-ter from Kir6.2 can reach and explore the sulfonylurea drug binding region in the middle of SUR1 and can keep SUR1 in the inward open form^17^. The disordered N-ter region from aa 1 to 32 is unsolved in most of the available structures with one exception of the fragment 1-20 captured in low resolution by Martin et al.^9^ in the drug-binding cavity of SUR1 being in the inward open state. The possible positions and localization of the N-ter when SUR1 is in the outward open conformation remain unknown. Numerous experimental data indicate the importance of N-ter in proper KATP functioning^9, 18–20^. Moreover, numerous mutations can disrupt its function, for example: L2P, S3C, A28-R32 deletion, F33I, R34C, and F35L cause NDM^21, 22^.

The C-terminal fragment of Kir6.2 (**C-ter**) from aa 353 to 390, absent in available structures, contains an endoplasmic reticulum retention motif (RKR) that prevents the cell’s surface expression of Kir6.2 in the absence of SUR1 subunit^23^. The RKR motif is in the region aa 369-371 and it is masked when SUR1 is present. Truncation of the gene encoding the last 26-36 amino acids of Kir6.2 causes expression of channels composed of Kir6.2 only^24^. The mutation of glycine 366 lying near the RKR motif to the much bulkier tryptophan causes NDM^25^.

The so-called lasso motif located between TMD0 and TMD1 (aa 192-286) (**L0-loop**) draws attention primarily by the abundance of pathogenic mutations (NDM related) occurring in this region^22^. Most of those mutations affect a fragment without a clear secondary structure - the N-terminal part of the L0-loop: I196N, P207S, E208K, D209N/E, Q211K, D212I/N/E/Y, L213R/P(DEND), G214R (MODY), V215I, R216C. In another protein of the ABCC family-CFTR, this motif provides a proper conformational stability of the core ABC structure by association with the membrane^26^. In KATP, the L0-loop may have an additional function. We know that the N-terminal L0-loop residues (199-214) contribute to the binding site of ATP in Kir6.2 (the so-called ABLOS-“ATP Binding Loop on SUR”)^27^. The ATP binding site partially overlaps with the PIP2 (phosphatidylinositol 4,5-bisphosphate) binding site^28^. PIP2 is a minor phospholipid from the inner leaflet of the membrane, which binding promotes the open structure of Kir6.2^29^. By the fact that the L0-loop directly connects the ABC core to an ATP/PIP2 binding site relevant in the opening-closing of the channel pore, L0-loop likely is one of the factors carrying the dimerization information of SUR1 domains into the channel pore^12^.

The spatial arrangement of the other SUR1 loops, i.e. **600-loop**, **700-loop**, and **900-loop**, places them in a position where they can significantly impact the dynamics of the N-ter and ABC domain of SUR1.

An un-modeled region extending from aa 620 to 677 of the SUR1 protein (**600-loop**) links the TMD1 and NBD1 domains. This fragment also contains the endoplasmic reticulum retention motif (RKR aa 648-650), preventing protein expression without KATP assembly. Similarly to C-ter, mutations near the RKR motif (R653Q) lead to transient neonatal diabetes^30^. KATP channels found in different human tissues contain different Kir/SUR subunits combinations. In pancreatic beta cells, these are Kir6.2 and SUR1, but in vascular smooth muscles, the latter is replaced by SUR2B. The L1 linker, a 600-loop analogue in SUR2B, has a phosphorylation site. Phosphorylation of the L1 linker activates the cardiac KATP channel^31^. No such activity appears in SUR1. Additionally, L1 in cardiac SUR2A has a heme-binding motif, absent in SUR1^32^. Thus, it could be that the additional (beyond RKR) functionality of the 600-loop in SUR1 has disappeared during evolution.

**700-loop** is a part of the NBD1 domain. It includes residues from 740 to 766. There is no information about possible posttranslational modifications of this region in the available literature. However, it has been reported that the nonsense mutation E747X causes neonatal diabetes^33^.

The undefined fragment ranging from aa 917 to 996 (**900-loop**) is intriguing as it contains the region especially rich in glutamic acid residues (the **ED domain**, following Sung et al.^10^). Glutamic acid is the second of the most common disorder-promoting residues^34^. Moreover, such a motif of seven glutamic acids in a row (aa 969-975), is conserved in all SUR-type proteins (SUR1, SUR2A and SUR2B) in all organisms. Interestingly, this motif does not appear in other ABC transporters, raising the possibility that it may be closely associated with the functioning of KATP channels. Indeed, the tripartite interaction between 900-loop, SUR2B and Kir6.1 has been recently reported for smooth muscle KATP channels^10^. It is known that glutamic acid/aspartic acid clusters coordinate various metal ions binding or destabilize the structure leading to weakening the ion binding capacity^34^. A reported mutation in this region causing neonatal diabetes that disrupts the channel function is R993C^30^.

### 2.2. Whole KATP system molecular dynamics simulations

In our relatively long simulation time (10 repetitions of 0.5 us each) the model of the system as a whole remained very stable. The characteristic propeller shape was retained, and SUR1 remained in the outward open conformation with NBD1 and NBD2 domains remaining close to each other and with MgADP bound at both consensus and degenerate binding sites (NBD1and NBD2). Fluctuations of the whole system are shown in SI in Figure S1. Selected ID regions had fluctuations reaching 5-12 Å above the general fluctuation level. The separate RMSDs of the fragments studied for each of the simulations performed (10 repeats x 4 protein chains) are shown in supplementary Figure S2.

Figure 2a shows the changes in the secondary structure of the disordered parts of the system during the simulation. In the case of the N-ter, we see a propensity for the α-helix formation for residues 8-16 and 20-30, but these are not stable structures. Above residuum 37, the stability of the structure is affected by several interactions with CTD. For C-ter, the disordered fragment starts around residuum 360 that ends the last ordered fragment of the CTD. Some fragments, such as aa 365-370, can form temporary α-helices, but those are short-lived phenomena. The L0-loop contains two helices, found in CryoEM. As expected, these fragments maintain the helical structure throughout the simulation. Among those helices, the structure is disordered and, apart from a slight tendency towards helicity for a fragment around residue 210, no significant regularities can be seen in the diagram. The other SUR1 loop analyzed, the 600-loop, shows the greatest disorder. The sporadic structures appearing are very short-lived and rather do not indicate specific interactions with the environment. In the last analyzed loop of SUR1, the 900-loop, the disordered fragment starts from residue 930, at the point where the α-helix belonging to NBD1 ends. In the diagram, we see the final fragment of this helix remaining in a stable form throughout the simulation. A further disordered fragment shows some tendency to helicity in the vicinity of residues 942 and 985.

**Figure 2.**
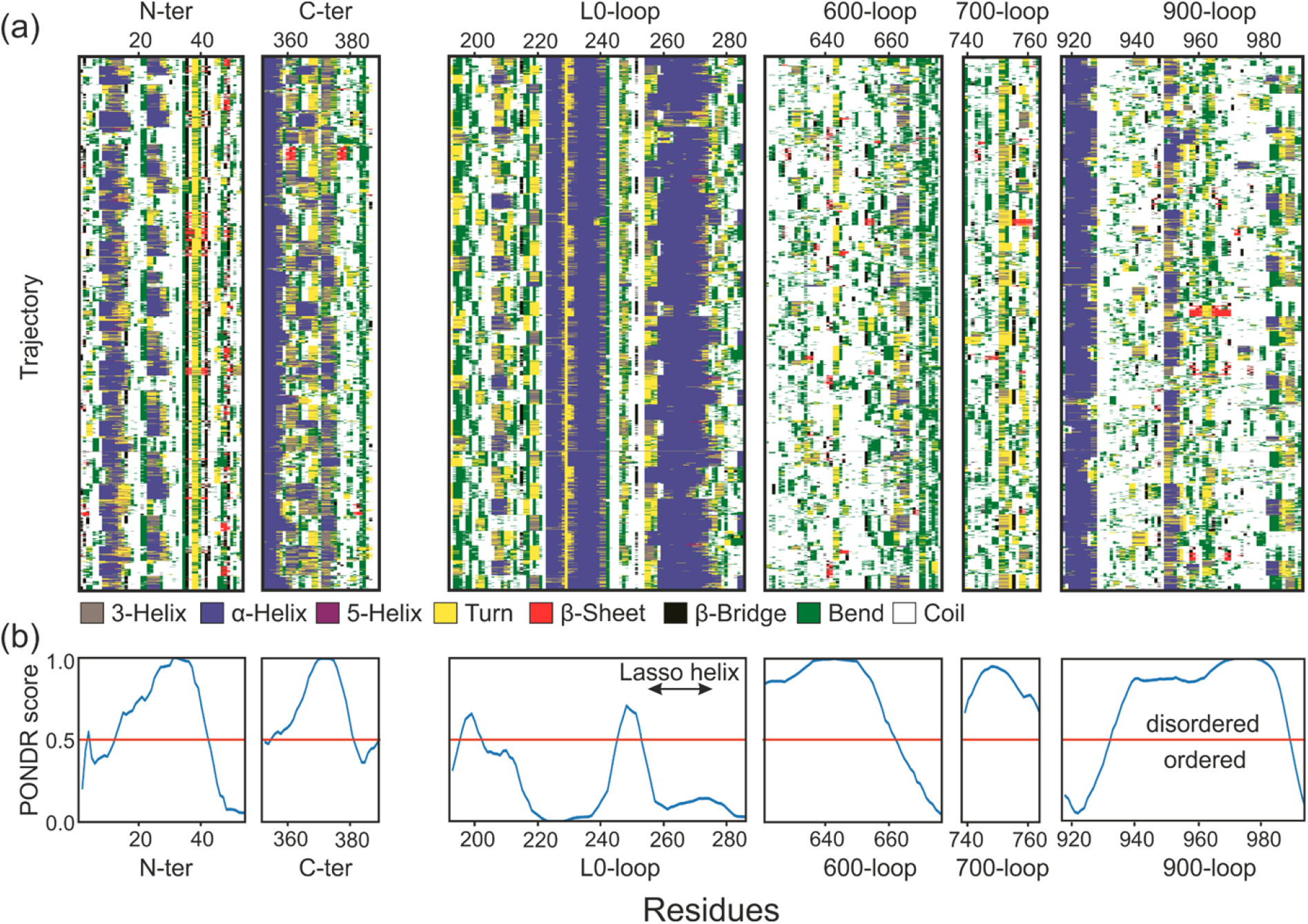
Time evolution of the secondary structure of IDRs as calculated by DSSP (a). The x-axis displays data collected during 0.5 μs for each chain, for 9 trajectories. Disorder in selected regions (b). The PONDR prediction tools were used to determine the disorder score. Any value above 0.5 indicates intrinsic disorder.

The disorder score calculated from the amino acid sequence reflects the dynamics of the fragments during the simulation. In particular, for N-ter, L0-loop, and 900-loop, where we also included fragments of ordered regions of the protein in the analysis.

### 2.3. Conformational space of unbound N-ter

Experimental studies^9, 18, 19, 35^ and our previous numerical studies^17^ indicate that Kir6.2 N-ter has an important function in the communication between SUR1 and Kir6.2. An extended fragment protruding from the channel shaft and docked in the SU pocket of SUR1 allows, from one side, to keep the CTD domains of Kir activated and, from the other side, to stabilize the inward open conformation of the ABC core. The presence of Kir N-ter in the interior of the ABC-core also affects the possible binding of sulfonylureas^36^. Moreover, the C-terminal part of N-ter is known to participate in the ATP binding^13, 37^. However, we still do not have any information on what happens with N-ter when it is not bound inside the SUR1 niche - that is, when the SUR1 conformation changes to outward open.

The performed MD simulations gave us information about the evolution of this fragment over time. Ten repetitions of a 0.5 μs simulation of a system containing four copies of the Kir protein provides us with a total of 20 μs sampling of the N-ter conformational space itself. Despite smaller than expected RMSD values oscillating between 2 and 6 Å (see supplementary Figure S2), the fragment does not appear to have a well-defined “parking place” in the overall KATP structure. N-ter moves relatively freely in the confined space between the CTD and the ABC-core (supplementary Figure S3) without entering into more permanent interactions with any part of the system - none of the traced contacts between residues forms for more than 50% of the simulation time (see Figure 3 c). The most frequent interactions (reaching 40%) are salt bridges formed between charged residues in the middle and end part of N-ter and residues from NBD1 of SUR1 (E19-K795 and R27-E784) and CTD of the same (E19-H276) or neighboring Kir6.2 (R25-E321). Interactions of N-terminal parts of N-ter with disordered 900-loop (R4-T949 and K5-T949) are also quite frequent, at the 30% level. This is, in our opinion, an interesting observation. The KATP system has such architecture and aa composition in this region that any permanent binding of N-ter is of low probability. It means that flexible N-ter is “ready” all the time to sneak into SUR1 and to stabilize open forms of SUR1 (channel is closed). Perhaps, such lack of permanent “docking positions” for N-ter facilities its role as a conformational lock (or “selector”). Tightly bound N-ter might be useless in this task, or its efficiency would be lower.

**Figure 3.**
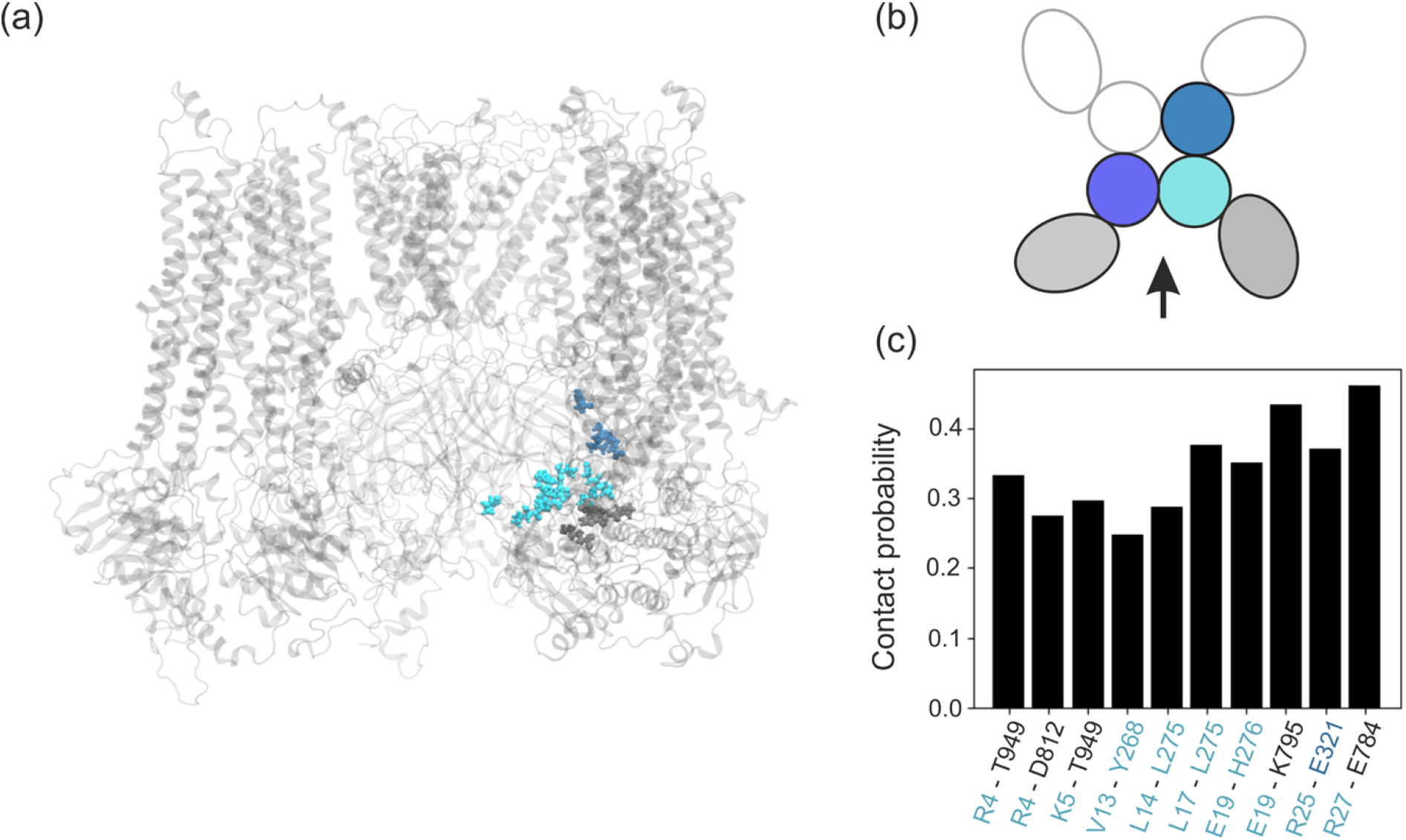
The spatial distribution of the residues most often in contact with N-ter assigned by color to the parts of the system to which they belong (a). Schematic, color-coded, representation of the KATP system. Colors are assigned to each part throughout the paper : shades of blue - individual Kir6.2 chains, gray and black -SUR1 (b). Frequency of close contact between N-ter residues and the rest of KATP (c).

### 2.4. L0-loop

L0-loop is not strictly disordered region. It has no defined structure in only one of the discussed conformations of the KATP - a quatrefoil state. Thus, it is almost completely well-determined by the available CryoEM structures in the propeller conformation. However, this region still has some flexibility, especially in its N-terminal side, whose amino-acid composition favors a lack of order (see PONDR score, Figure 2b). Although the space available for L0-loop movement is severely limited by the presence of other subunits of the system, the RMSD of the fragment in our simulations is of the order of 6 Å (supplementary Figure S2), implying substantial conformational freedom.

The L0-loop behavior is determined by stable interactions between the short helices comprising L0 and the TMD1 domain. These interactions occur in more than 80% of the simulations (see Figure 4 a,b). Of particular interest is that L0-loop directly interacts with the residues responsible for binding sulfonylureas (anti-diabetic drugs used to close the KATP channel), namely R1245 and E1246^36^.

**Figure 4.**
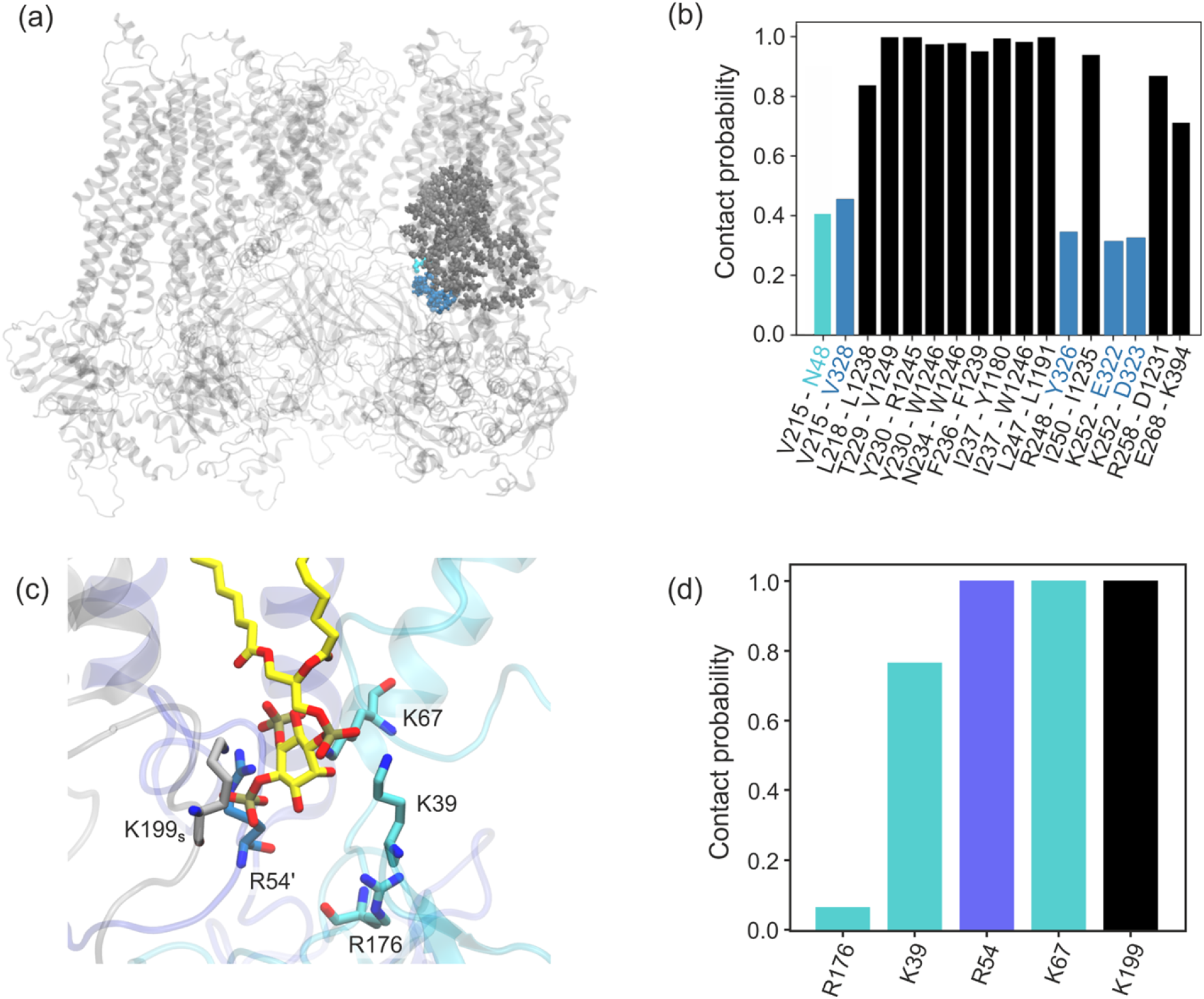
Spatial distribution of the residues most often in contact with L0-loop (a). Frequency of close contact between L0-loop residues and the rest of KATP (b). Close-up view of the Kir6.2 PIP2 binding site (c). Frequency of close contact between PIP2 and KATP residues (d).

In addition, experimental data indicate a direct involvement of the N-terminal part of L0-loop in ATP binding at the inhibitory site in Kir6.2. The CryoEM structure of Ding et al.^8^ shows that the side chain of K205 in SUR1 coordinates the β- and γ-phosphates of ATP. Therefore, these authors named residues 199-214 of SUR1 the “ATP Binding Loop on SUR’’ (ABLOS). Such motif is highly conserved from fish to humans^8^, and as mentioned before, several reported mutations in this region lead to NDM.

Our channel model is constructed to replicate the KATP open conformation as closely as possible: SUR1 is in the outward open conformation, and ATP at the inhibitory site of Kir6.2 is absent. Thus, we cannot explicitly show how the L0-loop participates in ATP binding. We can show that the L0-loop remains in direct contact (almost 40% of the simulation time) with the CTD domain of the neighboring Kir6.2 and N48, which is an extension of the N-ter region of the adjacent Kir6.2. The last residue N48, participates directly in ATP binding.

The absence of ATP at the inhibitory site of Kir6.2 and the presence of PIP2 in the lower membrane leaflet promotes PIP2 binding to KATP system. In our model, of the four available binding sites, in two PIP2 is stably bound to the channel, in one it is occasionally bound, and one site was vacant throughout all simulation (see SI). Our characterization of the PIP2 binding site is in agreements with recent numericall results^28^ (see Figure 4 c, d). It mainly consists of residues K39 and K67, and R54 from the neighboring Kir6.2 unit. The arginine R176, due to the rather strict definition of close contact adopted by us, is not directly involved in PIP2 binding, although it remains nearby. An exciting observation, not reported so far due to previous models’ limitations, is the direct participation of SUR1 in the PIP2 binding. The N-terminal L0-loop (ABLOS) lysine K199 (from SUR1) is one of the four positively charged amino acids that form the optimal environment for PIP2 binding.

### 2.5. Intrinsically disordered loops of SUR1 as a part of hypothetical molecular bearing

The flexible loops connecting NBD1 to the transmembrane domain are the most extended fully disordered fragments in KATP. Sterically, they are the only fragments of SUR1 subunits that can directly interact with adjacent ones. Therefore, one of the proposed functions performed by these fragments is to stabilize the characteristic propeller shape of the KATP complex. Indeed, contacts between neighboring SUR1s subunits through 600-loop and 900-loop loops occur in 20% of the simulation time (see Figure 5d). Another, KATP structure observed in CryoEM is quatrefoil conformation where SUR1 units are rotated counter-clockwise^12^. Nevertheless, since we have not considered the less frequent quatrefoil conformation of pancreatic KATP or the conversion between these two forms, such a hypothesis still needs to be confirmed. On the other hand, 600-loop and 900-loop provide a necessary flexibility (or conformational freedom) to all four SUR1 domain: the **600-loop** connects TMD1 to NBD1, the **900-loop** connects NBD1 to TMD2. At the same time presence of those ID loops probably reduce friction between Kir6.2 and SUR1 subunits during open/close KATP cycling. Thus, a critical analysis of dynamics of this IDR is presented here.

**Figure 5.**
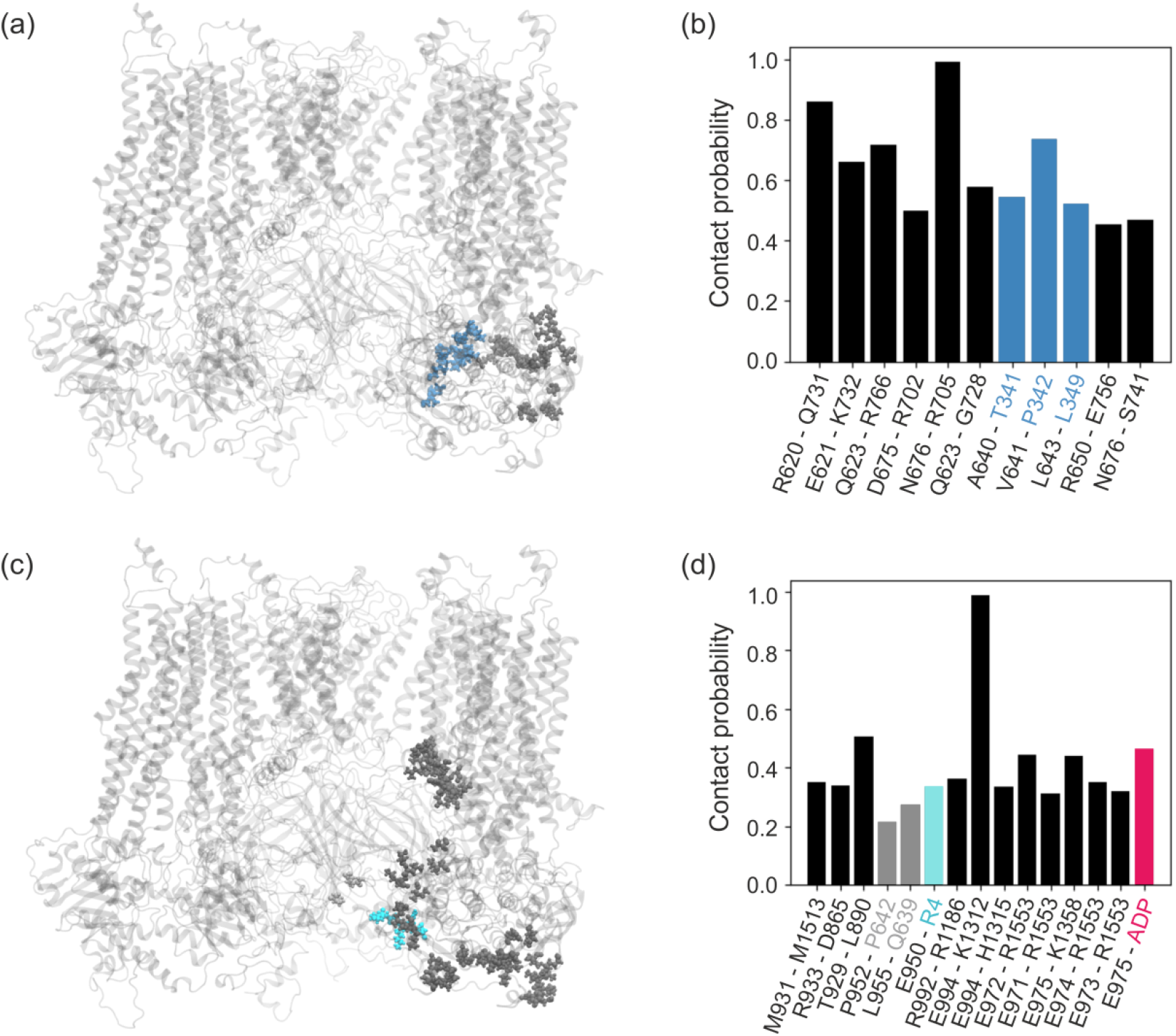
Spatial distribution of the residues being most often in contact with 600-loop (a) and 900-loop (c). Frequency of close contact between Kir6.2 and SUR1 residues and 600-loop (b), and 900-loop (b, d).

In SUR proteins (i.e., SUR1, SUR2A and SUR2B), the 600-loop is longer than in other proteins of the ABCC family (e.g. CFTR and MRP1 with known atomic structure), which may indicate some particular function this loop has in KATP channels. We have already mentioned that SUR1 has an endoplasmic reticulum retention motif, ensuring co-expression of SUR1 with Kir6.2. In SUR2B, on the other hand, it has been shown that 600-loop’s phosphorylation regulates channel activity^15, 38^ - disruption of the loop interactions with NBD1, leads to increased affinity of NBD1 for MgATP. NMR studies of Sooklal et al.^15^ on an isolated 600-loop - NBD1 system from rodent SUR2B show that the 600-loop possesses a residual secondary structure that is disrupted with phosphorylation. Such disruption leads to changes in the interaction pattern of the phosphorylated 600-loop with NBD1.

Despite the similar length of the 600-loop in SUR1, neither available experimental data nor numerical predictions indicate the existence of phosphorylation sites here (see UniprotKB database). Furthermore, if we assume some similarity in the functioning of the 600-loop with SUR1 and SUR2B, we would expect to observe some temporary structures in the un-phosphorylated state, stabilizing the 600-loop interactions with NBD1. The molecular dynamics data obtained for 600-loop of SUR1 do not show the formation of such transient structures (see Figure 2a). Nevertheless, the close contacts frequency analysis reveals a relatively stable interaction of 600-loop with NBD1 residues (Figure 5 a, b). Despite the large conformational freedom displayed by 600-loop (supplementary Figure S3), some of the resulting interactions are very stable, e.g. the pair of N676-R705 residues remains nearby almost throughout the simulation (see Figure 5b). Also, we observe frequent proximity for the residues R620-Q731, a particularly important feature as Q731 interacts directly with W688, which is a part of the ATP binding site in NBD1 (degenerate site). Additionally, the 600-loop residues interact with CTD of Kir6.2 adjacent to the unit interacting with SUR1 through the TMD0 domain. High flexibility of this CTD neighborhood facilitates functional motions of Kir6.2 CTD (molecular bearing). Due to the proximity in space of the 600-loop and 700-loop, interactions between them are also frequent.

The second fragment connecting NBD1 to the transmembrane part (TMD2) is the longest disordered fragment of SUR1. So far, little is known about its function. In other proteins of the ABCC family, i.e. CFTR and MDR1, this fragment is even longer (200 and 90 amino acids, respectively). In the former case, it forms a separate R domain, which regulates the protein’s function. Additionally, in both cases, it is phosphorylated which may modify the function of the fragment.

In SUR proteins, the 900-loop fragment is approximately 80 amino acids long. It does not contain experimentally confirmed phosphorylation sites, although prediction servers such as PhosphoSite suggest possible phosphorylation of serine 979 in SUR1. Due to its location at the system’s periphery, the 900-loop samples a significant volume over time, compared to other IDRs (see supplementary Figure S4). The PCA analysis allows us to distinguish three groups of the most common conformations, with different courses of the loop. There is a substantial anisotropy in the dynamics of the loop for each SUR1 subunit. Nevertheless, we can identify several hot spots in the interaction of the fragment with the rest of SUR1 (see Figure 5c,d). More than 40% probability of close contacts is shown by the pairs T929-L890 of NBD1 and E972-R1553 and E974-K1358. The first residue in the two last pairs belongs to an unusual accumulation of seven glutamic acids (ED), while the other belongs to the C-terminal and N-terminal parts of NBD2, respectively. The end of the disordered 900-loop fragment is determined by the highly stable salt bridge connecting E994 to K1312 belonging to TMD2. Residues 990-996 are also involved in the interaction with the membrane. Interestingly, the 900-loop region, especially the ED domain, interacts directly with the N-ter (almost 40% probability of contact with R4), and in more than 40% of cases, it co-forms the MgADP binding site (consensus site).

In Figure 6a, is shown the MgADP binding site, together with a fragment belonging to the ED domain of the 900-loop. Additionally, we analyzed the contact frequencies of amino acids lying in the vicinity of the ligand with the MgADP ligand itself (Figure 6b). The obtained results overlap to a large extent with the experimentally determined MgADP binding site for SUR1 in the outward open conformation^13^, i.e. R1110, Y1353, S1386, and K1384, make significant contributions. In our simulation, this set is augmented by V1360, R1379 and T1380 from NBD2. S830 belonging to NBD1, a part of the experimentally determined MgADP binding site, shows a contact probability of 20% due to our strict definition of ‘close contact’ (see Methods). The presence of residues from the glutamic acid-rich part of the 900-loop directly in the MgADP binding site suggests that this region may have arisen during evolution (and is conserved) specifically to affect MgADP binding. Indeed, Sung et al. showed MgADP dependent interaction among 900-loop, Kir6.1, and SUR2B in smooth muscle KATP channels in the quatrefoil conformation^10^. They propose a model in which the glutamic-acid-rich part of 900-loop functions as a gatekeeper to prevent unregulated channel activation in the absence of MgADP. Nevertheless, the relevance of such a mechanism in the pancreatic KATP system is not obvious. It is mainly because the quatrefoil conformation, common in the vascular KATP, is not the dominant one in case of pancreatic KATP.

**Figure 6.**
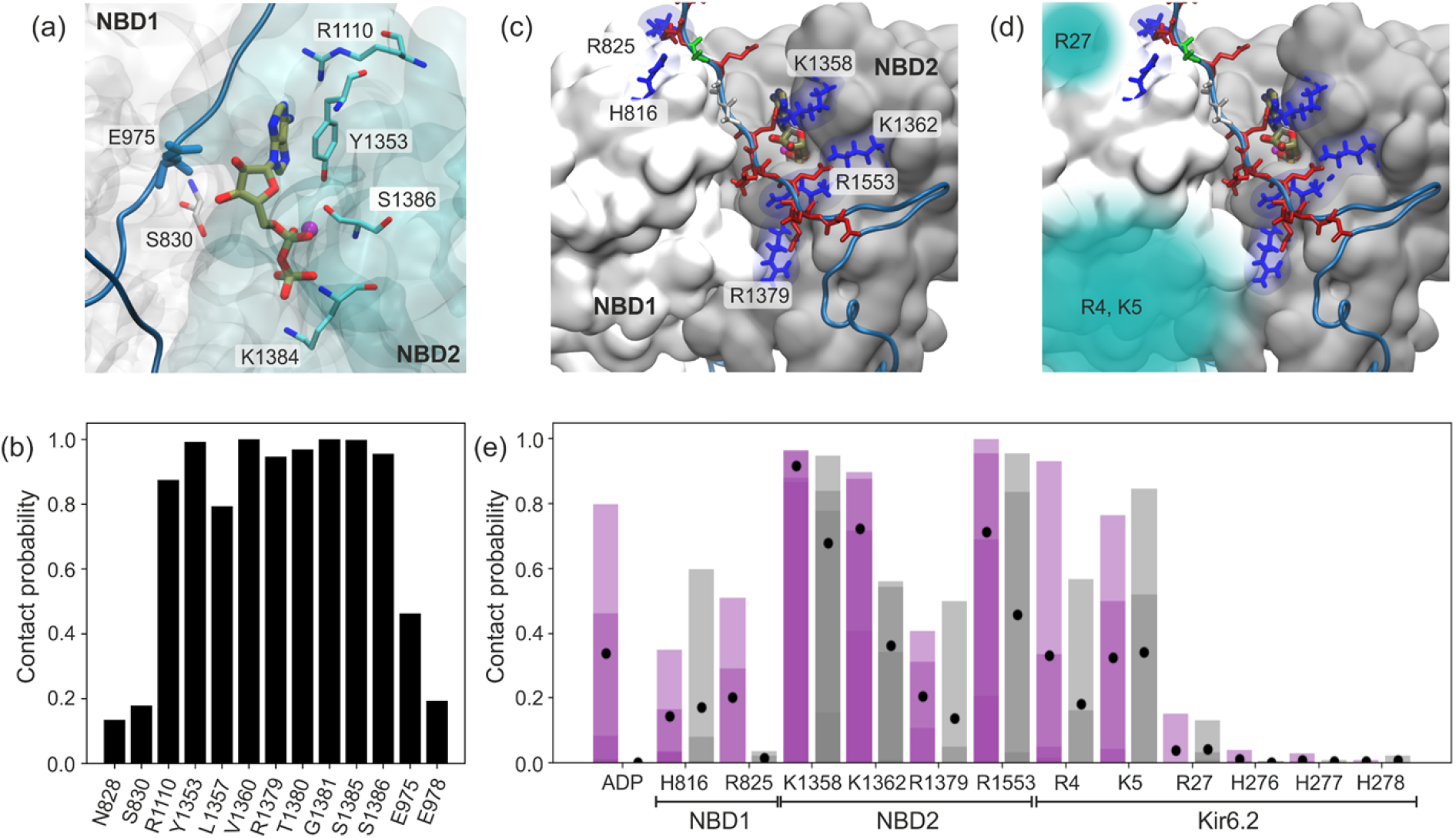
Close-up view of the consensus MgADP binding site in SUR1 NBDs (a). Frequency of close contact between MgADP and KATP residues (b). Location of positively charged anchors in NBD1 and NBD2 (blue), interacting with the ED domain (red) (c). Areas where interaction with the N-ter can occur (d). Frequency of contacts between ED domain and the “anchor” residues in two cases: ADP present in the binding pocket (magenta) and the APO model (gray) (e). Due to the high anisotropy between KATP units (chains), the contact frequencies of each chain are shown together as transparent bars. The averaged value is marked with a black dot.

To test how the presence of the ED domain in the ADP binding site affects ligand dynamics in pancreatic KATP, we examined its fluctuations over time. During the evolution of the system, the NBD domains forming the MgADP binding site spontaneously tighten and relax yet remain closed - the distance between the centers of mass of the NBD1-NBD2 domains varies from 28 to 36 Å, compared to 45 Å. in the inward open SUR1 conformation. The tight closure of the domains limits the mobility of ADP, which could potentially lead to its dissociation. The average RMSD for ADP in the tight domain closure (the distance between COM NBDs < 29.5 Å) is 1.84 Å (0.17). For such an arrangement of domains, the interaction of the ED domain with ADP occurs sporadically.

A slight relaxation of the NBD domains (COM distance >30.5 Å) changes the situation. Here, the ED domain has free access to the ligand and interaction between the adenosine ribose hydroxyl group and the backbone (mainly oxygen in the carboxyl group) of the E975 residue occurs very frequently (see Figure6 a, b). The frequency of this interaction depends mainly on the degree of closure of the NDB domains and shows high anisotropy for different units. Such a rather unusual interaction of negatively charged glutamic acid with ADP results mainly from the interaction of the additional carboxyl group of E976 with the neighboring lysine L1358 (see Figure 6 c). The presence of the ED domain seems to have a large stabilizing effect on the ADP binding in NDBs - the measured RMSD of the ligand for relaxed domains is 2.07 Å (0.11) in the absence of ED, and when the interaction with ED occurs, the RMSD drops to 1.75 Å (0.1). A smaller RMSD probably translates into a lower chance of ligand dissociation. However, this hypothesis requires further computational study.

The ED domain, naturally retaining some freedom due to its disordered nature, probably prefer this specific arrangement in order to limit ADP fluctuations, i.e., it anchors itself by interacting with positively charged residues of NBD domains at the interface.

One hypothesis proposed by Sung et al. is that the ED domain is a bridge between NBDs and CTDs. It may be involved in relaying the signal indicating the presence of ADP (and dimerization of NBD domains) in SUR2B directly to the CTDs in Kir6.1. We analyzed such a possibility for pancreatic KATP case. In our channel model, despite relatively long simulation times and a decent statistical sample, the NBD domains remain dimerized. We did not observe the diffusion of MgADP from the binding sites, even though relaxation of the NBDs domains sometimes leads to quite significant fluctuations of the ligand. Thus, to compare the structural dynamics of the l900 loop in the presence and absence of ligand, we used the APO model as a reference. In such a model, the MgADP binding sites in all NBDs are unoccupied. We analyzed the frequency of close contacts between the ED domain and most probable anchors in two cases: in the presence of ADP and without, each time for four different chains (see Figure 6e). Our data show that ED interacts very little with the CTD domain, making it unlikely that the signal is transmitted in that way. The contact with the Kir6.2 part of the system occurs mainly through contacts with the N-tail. Here, the positively charged R4 and K5 contribute the most (albeit with high chain related anisotropy), and R27 occasionally interacts with ED (see Figure 6d,e). There are sporadic interactions with the histidine series in the CTD. It is important to note that the percentage of ED interactions with anchors varies with the degree of closure of the NBD domains (supplemental Figure S5). Direct interactions with ADP barely occur for tightly closed domains, interactions with K1362, R1379 and R1553 become less frequent, and interactions with histidins in CTD are virtually nonexistent.

## 3. Conclusions

The role of IDRs of proteins in living organisms is becoming better understood. Considering a large number of such regions recognized so far^1, 39^, both cytosolic and membrane, we are probably only at the beginning of this route. In the case of the KATP channel, vital for the proper functioning of human body, our studies allow us to unveil the possible functional roles of yet unexplored fragments.

First of all, these regions may participate in “transferring information” between domains via modulation of subunit-subunit interactions. In particular, this refers to the N-ter Kir6.2 tail, which we described in our recent paper^17^, and the L0-loop described here. These fragments serve as links between SUR1, being an ATP intracellular concentration sensor, and the channel pore formed by Kir6.2 four subunits. Probably concerted motions in these two pathways lead to correct functioning of the complex.

Moreover, we show computationally that IDRs are involved in physiological ligands binding. The L0-loop, particularly the ABLOS part, contributes to the binding of both ATP and PIP2. The 900-loop, on the other hand, through the existence of a very rare glutamic acid-rich fragment, directly affects the MgADP binding site in NBD1 SUR1. Such stabilizing interaction prevents the excessive ligand fluctuations, which might otherwise lead to the ligand dissociation during spontaneous NBD domains relaxation.

Additionally, our study shows that the transient direct interactions between adjacent SUR1 units are possible through the 600-loops and 900-loops. Such contacts may modulate the stability of the characteristic propeller structure of the KATP channel. Due to location of 600-loop at the close proximity to CTD domain of Kir6.2 its high flexibility may be also functional – it may be a part of hypothetical of molecular bearing type Kir6.2/SUR1 cytosol located interface. Significance of this findings has yet to be confirmed by further investigations.

## 4. Methods

### 4.1. System preparation

The fully solvated KATP complex model was built based on the CryoEM structure of the human KATP in the activated form (PDB ID: 6C3P)^12^. Each Kir6.2 and SUR1 part was examined for missing side chains and loops using the Schrodinger software. The minor missing regions (intra and extracellular loops of SUR1 shorter than 20 aa) were filled in using Prime-Schrodinger module, whereas two major missing regions in SUR1 (600-loop, and 900-loop, see figure 1c) were modelled using the I-TASSER structure prediction server^40, 41^. The structure of the missing terminal parts of Kir6.2 subunits were predicted using the QUARK server^42^. The newly added fragments were pre-equilibrated as described in Walczewska-Szewc and Nowak, 2020^17^.

Using Charmm-gui server^43–46^, the model of KATP was embedded into an explicit palmitoyloleoyl-phosphatidylcholine (POPC) bilayer with 10% phosphatidylinositol 4,5-bisphosphate (PIP2) in the inner leaflet of the membrane^46^. Two MgADP molecules (with force-field parameters generated by Charmm-gui) were placed in consensus and degenerate binding sites in each SUR1. For the APO models, the binding sites remained unoccupied. The entire protein–membrane systems were solvated with the TIP3P water and 150 mM KCl, resulting in a simulation box 250×250×210 Å^3^ and consisted of approximately 1.3 mln atoms.

### 4.2. MD simulations

All MD simulations were performed with GROMACS 2019^47^. Bonds to hydrogen atoms were held rigid using LINCS algorithms, allowing us to use 2 fs time step. Periodic boundary conditions were used. Long-range electrostatic forces were calculated using the Particle Mesh Ewald method^48^. The simulations were performed with a constant temperature of 310 K maintained using Nose-Hoover thermostat^49^ and a constant pressure of 1 atm (Parrinello-Rahman algorithm^50^). The CHARMM36 force field^51, 52^ was used in all the MD simulations. After minimization and equilibration period (20 ns), ten replicas of the system were run. Each simulation gave us a 500 ns trajectory. Two additional 500 ns simulations were run for the APO model.

### 4.3. Data analysis

The structure assignment for each residue and time was calculated using Gromacs *do_dssp* tool. After reading a trajectory file, it computes the secondary structure for each time frame calling the *dssp* program^53^. The PONDR score for each region was calculated using the Predictor of Natural Disordered Regions server (pondr.com).

Frequency of a ligand-residue and residue-residue contacts were calculated using residueresidue contact score^54^ for each frame across the trajectory, setting the 0.1 threshold. RMSD, RMSF, PCA and all structural properties of the system were calculated using Python scripts that utilize *MDAnalysis*, *MDTraj*, *NumPy*, *SciPy*, *scikit-learn* and *Matplotlib* libraries. Couplings among parts of PREP in each system were analyzed using the dynamic crosscorrelation (DCC) matrices^55^. VMD was used for molecular structures visualization^56^.

## Acknowledgements

The authors acknowledge funding from the National Science Centre, Poland (Grant 2016/23/B/ST4/01770). The computational results used in this paper were obtained with the support of the Interdisciplinary Centre for Mathematical and Computational Modelling ICM, University of Warsaw, under Grant GA76-10 and the facilities of the Interdisciplinary Centre for Modern Technologies, NCU, Poland. KWS acknowledges the support from the NCU Centre of Excellence „Toward Personalized Medicine”.

